# ATML1 Regulates the Differentiation of ER Body-containing Large Pavement Cells in Rosette Leaves of Brassicaceae Plants

**DOI:** 10.1101/2023.07.28.551031

**Authors:** Alwine Wilkens, Paweł Czerniawski, Paweł Bednarek, Marta Libik-Konieczny, Kenji Yamada

## Abstract

Endoplasmic reticulum (ER)-derived organelles, ER bodies, participate in the defense against herbivores in Brassicaceae plants. ER bodies accumulate β-glucosidases, which hydrolyse specialized thioglucosides known as glucosinolates to generate bioactive substances. In *Arabidopsis thaliana*, the leaf ER (LER) bodies are formed in large pavement cells, which are found in the petioles, margins, and blades of rosette leaves. However, the regulatory mechanisms involved in establishing large pavement cells are unknown. Here, we show that the ARABIDOPSIS THALIANA MERISTEM L1 LAYER (ATML1) transcription factor regulates the formation of LER bodies in large pavement cells of rosette leaves. Overexpression of *ATML1* enhanced the expression of LER body-related genes and the number of LER body-containing large pavement cells, whereas its knockout resulted in opposite effects. ATML1 enhances endoreduplication and cell size through LOSS OF GIANT CELLS FROM ORGANS (LGO). Although the overexpression and knockout of *LGO* affected the appearance of large pavement cells in Arabidopsis, the effect on LER body-related gene expression and LER body formation was weak. LER body-containing large pavement cells were also found in *Eutrema salsugineum*, another Brassicaceae species. Our results demonstrate that ATML1 establishes large pavement cells to induce LER body formation in Brassicaceae plants, contributing to the defense against herbivores.

## Introduction

Plants are equipped with a wide variety of physical and chemical defense mechanisms to protect themselves from pathogens and pests (Gong and Zhang, 2014). The Brassicaceae family plants have evolved the so-called “mustard-oil bomb system” as a chemical defense system (Lüthy and Matile, 1984; Matsushima et al., 2002), which relies on the precursor form of repellent metabolites, called glucosinolates, and a group of thio-β-glucosidases (thio-BGLUs), called myrosinases. The enzymes and substrates are stored in different tissues or subcellular compartments, but damage to the tissues or cells, e.g., through herbivory, enables direct contact between the enzymes and substrates, leading to the deglycosylation of glucosinolates (Kissen et al., 2009) and subsequently the production of various metabolites, including toxic isothiocyanates. The canonical myrosinases are localized in unique cells, called myrosin cells, in the vasculature system (Kissen et al., 2009), and it was previously believed that those myrosinases are the only enzymes responsible for the activation of glucosinolates. However, recent studies show that BGLUs that localize to endoplasmic reticulum (ER) bodies (e.g., PYK10/BGLU23), also exhibit myrosinase (thio-BGLU) activity and are important for the resistance against herbivores (Yamada et al., 2020).

ER bodies have been described as spindle-shaped organelles, 5–10 µm in length, which are directly connected to the ER (Matsushima et al., 2003a, 2002). Depending on their expression site, regulatory pathways, and components, ER bodies can be classified into categories: constitutively expressed ER (cER) bodies, inducible ER (iER) bodies, and leaf ER (LER) bodies. The cER bodies are present in the epidermal cells of cotyledons, hypocotyls, and roots (Matsushima et al., 2003b). Accumulation of cER bodies is regulated through the NAI1 transcription factor, which binds to the promoter regions of *NAI2* and *PYK10* (Sarkar et al., 2020). NAI2 is an ER body-localized protein responsible for cER body formation (Yamada et al., 2008), while PYK10 is the major component of cER bodies. The iER bodies are formed in the epidermal cells of mature rosette leaves but only after wounding or treatment with the wounding-related phytohormone jasmonic acid (JA) (Matsushima et al., 2003a, 2002; Ogasawara et al., 2009). Together with NAI2, TONSOKU-ASSOCIATING PROTEIN 1 (TSA1), a homolog of NAI2, regulates the formation of iER bodies, whose main component is BGLU18, a homolog of PYK10. LER bodies are formed in the epidermal cells of rosette leaves but are restricted to large pavement cells, which are formed at leaf margins, in the petiole, and at random points throughout the leaf blade (Nakazaki et al., 2019a; Roeder et al., 2012). LER bodies contain NAI2 and PYK10 (cER body-related components) as well as BGLU18 (iER body-related component) (Nakazaki et al., 2019a). In rosette leaves, NAI1 regulates the accumulation of LER bodies by regulating *NAI2* and *PYK10* expression, while CORONATINE INSENSITIVE 1 (COI1), a JA receptor, regulates *BGLU18* expression (Nakazaki et al., 2019a, 2019b). Therefore, LER bodies combine several characteristics of cER and iER bodies (Nakazaki et al., 2019a; Ogasawara et al., 2009).

In the rosette leaves of *Arabidopsis thaliana*, large pavement cells are characterized by their physically large size, which is associated with endoreduplication (Melaragno, 1993), a specialized cell cycle (Sugimoto-Shirasu and Roberts, 2003; Tsukaya, 2019) that increases the cellular ploidy level (Larkins et al., 2001). Similarly, large endoreduplicated epidermal cells, termed giant cells (GCs) (Roeder et al., 2012), are formed in Arabidopsis sepals, and their gene expression profile is different from the gene expression profile of the surrounding smaller cells (Roeder et al., 2012). GCs in the sepals of Arabidopsis express high levels of many defense-related genes, including (L)ER body-related genes (Schwarz and Roeder, 2016). Therefore, it is likely that GCs in sepals and large pavement cells in rosette leaves constitute the same cell type. Several genes are associated with the formation of GCs in sepals, namely, *ARABIDOPSIS THALIANA MERISTEM L1 LAYER* (*ATML1*), *DEFECTIVE KERNEL1* (*DEK1*)*, ARABIDOPSIS CRINKLY4* (*ACR4*)*, HOMEODOMAIN GLABROUS11* (*HDG11*), and *LOSS OF GIANT CELLS FROM ORGANS* (*LGO*; also known as *SIAMESE RELATED 1* [*SMR1*]) (Roeder et al., 2012). Notably, *DEK1*, *ACR4*, and *HDG11* are involved in epidermal cell differentiation, but these genes are also responsible for the establishment of the GC-specific gene expression profile, hereafter referred to as GC identity (Roeder et al., 2012; Iida and Takada, 2021; Meyer et al., 2017). According to previous studies, establishment of the GC cell type is followed by the induction of *LGO*, which triggers endoreduplication but has no influence on gene expression (Roeder et al., 2012). However, the results of recent RNA-seq studies suggest that *LGO* also influences overall gene expression in GCs (Roeder et al., 2012; Schwarz and Roeder, 2016).

Although the key components of LER bodies have been described, the genetic regulation of LER body-related genes is not fully understood. Moreover, the relationship between large pavement cell differentiation and LER body formation remains unclear. Here, we show that *ATML1*, a gene regulating the formation of GCs in sepals, also regulates the differentiation of large pavement cells and is responsible for the formation of LER bodies in Arabidopsis rosette leaves. We have also demonstrated the presence of LER body-containing large pavement cells in *E. salsugineum*, proving the universality of this defense mechanism in the Brassicaceae family.

## Results

### ATML1 regulates LER body-related gene expression to a greater extent than LGO

Because large pavement cells in rosette leaves and GCs in sepals show close resemblance in terms of being large endoreduplicated epidermal cells, we hypothesized that genes regulating the formation of GCs in sepals might also be responsible for the differentiation of large pavement cells and consequently for the induction of LER body formation in rosette leaves. To validate our hypothesis, we employed *ATML1*– and *LGO*– aberrant expression (overexpression and knockout) lines. *ATML1* is a transcription factor gene that regulates GC differentiation, while *LGO* controls endoreduplication but not GC differentiation (Roeder et al., 2012). Therefore, the analysis of these lines enabled us to determine whether cell type, endoreduplication, or neither is responsible for the formation of LER bodies in endoreduplicated large pavement cells.

First, we determined the mRNA levels of LER body-related genes, namely *NAI1*, *NAI2*, *PYK10*, and *BGLU18*, in the rosette leaves of 14-d-old plants (Figure 1A). Quantitative reverse transcription polymerase chain reaction (qRT-PCR) revealed that *ATML1* had a strong impact on the expression of the investigated genes; the expression levels of all examined genes, except *BGLU18*, were significantly enhanced in the *ATML1*-overexpressing (*ATML1*-OX) line. The expression of *NAI1*, a transcription factor gene that regulates LER body formation, was more than four times as high as in the wild type (WT). By contrast, the mRNA levels of *NAI1* and *PYK10* were significantly reduced in the *atml1-3* knockout line. *LGO* seemed to have a limited effect on the expression of ER body-related genes; while the mRNA levels of LER body-related genes showed no significant change in the *lgo*-2 knockout line compared with the WT, the mRNA levels of *NAI1*, but not those of *NAI2*, *BGLU18*, or *PYK10*, were significantly enhanced in the *LGO*-overexpressing (*LGO*-OX) line. Immunoblotting using antibodies directed against NAI2, BGLU18, and PYK10 revealed a strong reduction or increase in the levels of all tested proteins in *atml1-3* or *ATML1*-OX plants, respectively (Figure 1B), consistent with the qRT-PCR results. Interestingly, *lgo-2* or *LGO*-OX also displayed a reduction or increase in the levels of the tested proteins, respectively. However, the changes in protein accumulation were much smaller in *LGO*-aberrant expression lines than in *ATML1*-aberrant expression lines. These results suggest that *ATML1* and *LGO* influence both the mRNA and protein levels of LER body-related genes. However, the influence of *ATML1* was much greater than that of *LGO*, indicating that ATML1 plays a major role in regulating LER body-related genes, while LGO-mediated endoreduplication has only a small effect on the expression of LER body-related genes and proteins.

**Figure 1:**
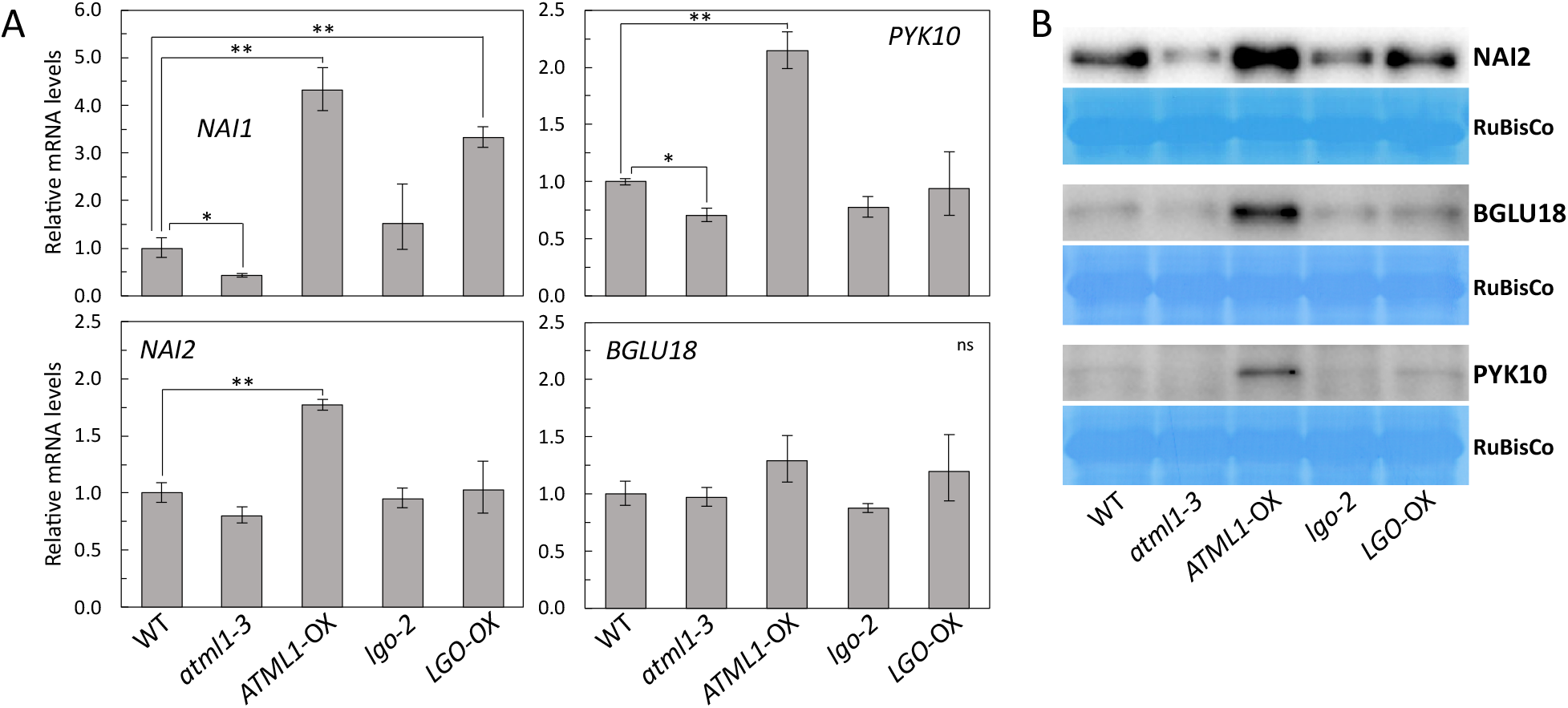
Giant cell (GC)-aberrant Arabidopsis lines show altered levels of LER body-related mRNAs and proteins in rosette leaves. **A**, Quantitative reverse transcription PCR (qRT-PCR) analysis of LER body-related genes in the rosette leaves of 14-d-old plants. Asterisks indicate significant differences compared with the wild type (WT; *p < 0.05, **p < 0.005; n = 3). **B**, Immunoblotanalysis of the first and second rosette leaves of 14-d-old plants using anti-NAI2, –BGLU18, and –PYK10 antibodies.

### ATML1 regulates LER body formation

In WT Arabidopsis rosette leaves, LER body-containing large pavement cells were abundant in petioles and leaf margins, and were less frequent in leaf blades (Nakazaki et al., 2019a) (Figure 2A). Because large pavement cells in petioles are easy to monitor, we examined the formation of LER bodies using the petioles of *ATML1*– and *LGO*-aberrant expression lines that stably expressed ER-localized GFP (GFP-HDEL) (Haseloff et al., 1997). In the WT, large pavement cells with LER bodies covered large parts, but not the entirety, of the petiole (Figure 2B). Compared with the WT, LER bodies detected in the *atml1*-*3* knockout line were mostly smaller and were present in only a few cells (Figure 2B). Furthermore, the size of LER body-containing cells was reduced in *atml1*-*3*, suggesting the reduction of endoreduplication levels (Melaragno, 1993). On the contrary, petioles in the *ATML1*-OX line showed increased numbers of LER body-containing large pavement cells. Interestingly, aberrant expression of *LGO* greatly influenced the size of epidermal cells but had little effect on the number of LER bodies. In the *lgo-2* knockout line, epidermal cells were greatly reduced in size, which is a consequence of the reduced endoreduplication levels (Hamdoun et al., 2016), but several small cells harbored LER bodies. By contrast, in the *LGO*-OX line, the number of large pavement cells was increased, but several of these cells did not harbor LER bodies. To quantify the effect of *ATML1*– and *LGO*-aberrant expression on LER body formation, we calculated the percentage of the petiole area covered by epidermal cells that contain LER bodies while excluding stomatal cells from the measured area (Figure 2C). This analysis revealed that overexpression of *ATML1* increased the area of LER body-containing cells to ∼53%, while knockout of *ATML1* decreased the area to ∼14%, compared with the WT (∼27%). By contrast, the area of LER body-containing cells did not change in the *LGO*-aberrant expression lines compared with the WT. These findings indicate that *ATML1* regulates LER body accumulation as well as large pavement cell differentiation, and suggest that *LGO*-mediated endoreduplication and cell enlargement are dispensable for LER body accumulation.

**Figure 2:**
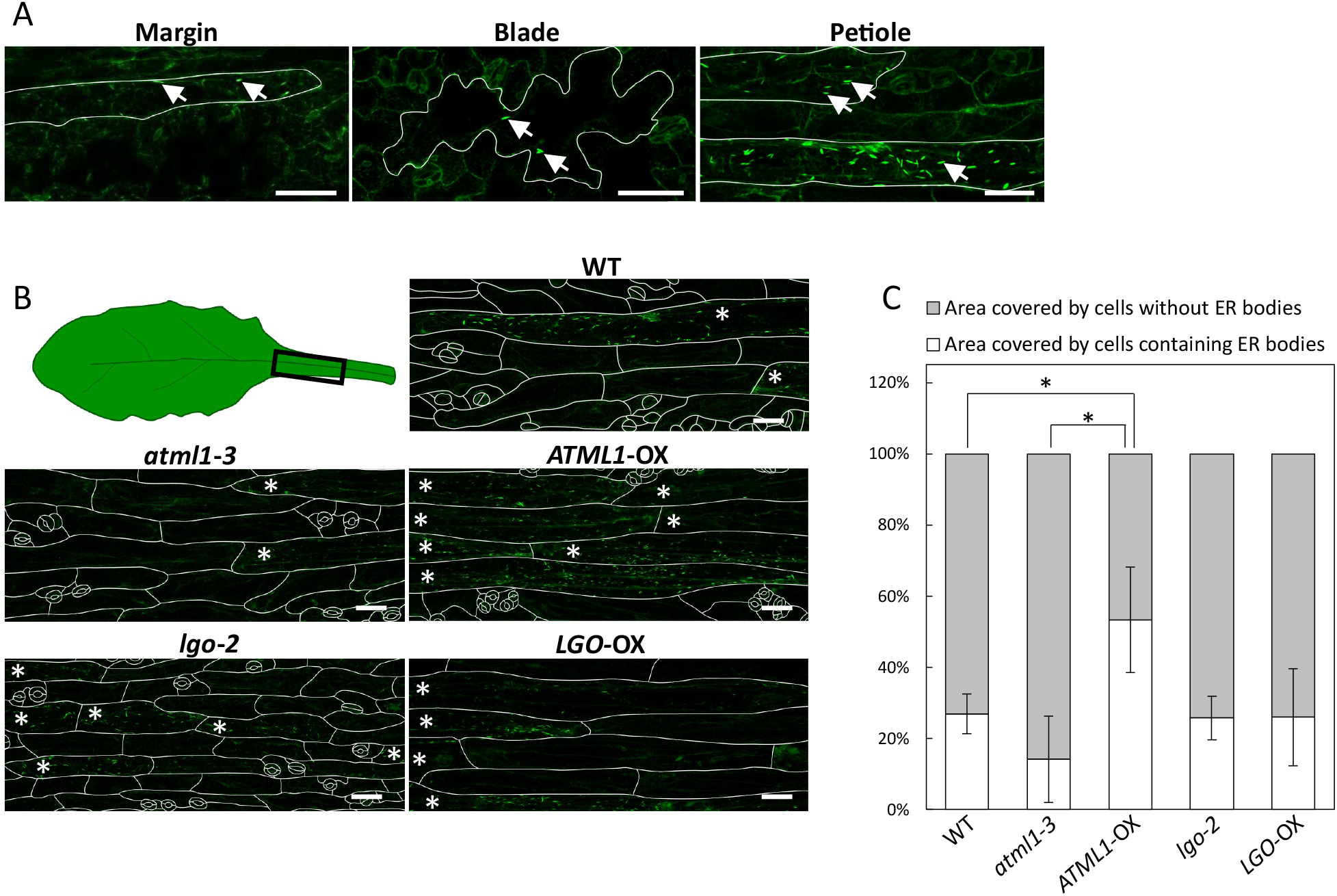
Influence of *ATML1* and *LGO* on the formation of LER bodies in Arabidopsis. **A**, LER bodies in the large pavement cells at the leaf margin and in the leaf blade and petiole. The boundaries of LER body-containing large pavement cells are outlined in white. Exemplary LER bodies are marked with arrows. Scale bars: 50 µm. **B**, LER bodies in *ATML1*– and *LGO*-aberrant expression lines. The epidermal cells of the petiole of 14-d-old plants were photographed using a camera-equipped confocal laser scanning microscope. All genotypes constitutively expressed ER-localized GFP-HDEL, and ER bodies were detected via the green channel. Cell sizes as well as ER body numbers varied greatly among the different Arabidopsis lines. Cells containing LER bodies are marked with asterisks, and cell boundaries are outlined in white. Scale bars: 50 µm. **C**, Percentage of the petiole area covered with epidermal cells containing (white) or lacking (grey) LER bodies in Col-0 (WT), *atml1-3*, *ATML1*-OX, *lgo-2*, and *LGO*-OX plants. Stomata were excluded from the measurements. Five pictures were analyzed per genotype. Asterisks indicate significant differences compared with the WT (*p < 0.05; n = 3).

Because LER bodies are involved in chemical defense using glucosinolates, we examined the glucosinolate content of the rosette leaves of *ATML1*– and *LGO*-aberrant expression lines. Compared to the WT, *atml1*-*3* showed a clear and significant enhancement of glucosinolates, which is possibly due to the reduction of PYK10 (Supplemental Figure 1, Supplemental Table 1). However, *lgo-2* and *LGO*-OX also displayed significant increases in glucosinolate levels.

### ATML1 regulates *NAI2* and *PYK10* expression in an indirect manner

To further investigate if the regulation of LER body-related genes by ATML1 is NAI1– dependent, we measured *NAI2*, *PYK10*, and *BGLU18* mRNA levels in the rosette leaves of WT, *nai1-1*, *ATML1*-OX, and *nai1-1*/*ATML1*-OX plants (Figure 3). In the *nai1-1* mutant, *NAI2* and *PYK10* mRNA levels were significantly reduced, while *BGLU18* mRNA levels were slightly enhanced. This suggests that NAI1 regulates *NAI2* and *PYK10* expression, whereas *BGLU18* expression is independent of NAI1. Overexpression of *ATML1* significantly increased the mRNA level of all tested genes; however, knockout mutation of *NAI1* in the *ATML1-*OX line decreased *NAI2* and *PYK10* mRNA levels to those observed in the *nai1-1* knockout line. Therefore, expression levels of *NAI2* and *PYK10* in *ATML1*-OX are dependent on NAI1. On the contrary, *BGLU18* mRNA levels did not decrease in the *nai1-1*/*ATML1*-OX line compared with the WT, demonstrating that the induction of *BGLU18* is not dependent on NAI1 in the *ATML1*-OX line.

**Figure 3:**
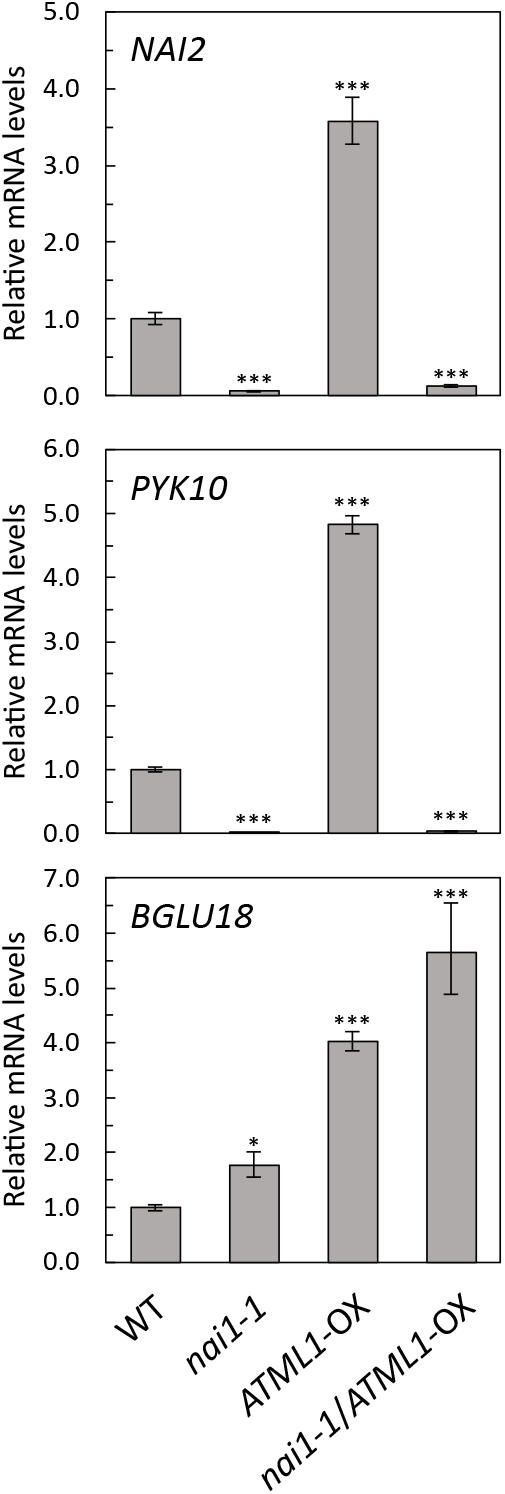
Influence of *nai1* mutation on ATML1-dependent LER body-related gene expression. Relative mRNA levels of *NAI2* (top), *YK10* (middle), and *BGLU18* (bottom) in the rosette leaves of Col-0 (WT), *nai1-1*, *ATML1*-OX, and *nai1-1*/*ATML1*-OX plants. Asterisks dicate significant differences compared with the WT (*p < 0.05, ***p < 0.0005; n = 3).

ATML1 activates gene expression by binding to the so-called L1 box, an 8 bp cis regulatory element (5’-TAAATG(C/T)A-3’) (Abe et al., 2001). We searched for the L1 box motif in the promoter regions (1 kb sequence upstream of the transcription start site) of *NAI1*, *NAI2*, *PYK10*, and *BGLU18*. We found two L1 boxes in the *NAI1* promoter and one L1 box in the *BGLU18* promoter (Supplemental Figure 2A, B), suggesting that these genes are directly regulated by ATML1. Congruently, the levels of *BGLU18* mRNA as well as its protein were increased not only in rosette leaves (Figure 1) but also in cotyledons of the *ATML1*-OX line (Supplemental Figure 3). No L1 box could be detected in *NAI2* and *PYK10* promoter regions; however, they contained G– and E-boxes, which are known to serve as NAI1 binding sites (Supplemental Figure 2C, D) (Nakazaki et al., 2019a). Therefore, our findings suggest that ATML1 regulates *NAI1* and *BGLU18* in a direct manner but regulates *NAI2* and *PYK10* indirectly by enhancing NAI1 levels.

### LER bodies are formed in Eutrema salsugineum

To determine whether Brassicaceae plants other than Arabidopsis harbor LER bodies, we examined LER body accumulation in *E. salsugineum,* whose genome sequence is publicly available (Batelli et al., 2014). We stably transformed *E. salsugineum* plants with *GFP-HDEL* to visualize the ER and ER bodies. Large pavement cells containing LER bodies were detected in GFP-HDEL-expressing *E. salsugineum* plants (Figure 4), specifically at the margin and in the petiole and blade of rosette leaves, as observed in Arabidopsis. However, LER bodies in *E. salsugineum* were longer and not as straight as those in Arabidopsis. Interestingly, we found that the promoter region of the *NAI1* homolog in *E. salsugineum* also contained an L1 box (Supplemental Figure 2E), which suggests that ATML1 functions similarly in both *E. salsugineum* and Arabidopsis, i.e., upregulates *NAI1* expression to induce the accumulation of LER bodies.

**Figure 4:**
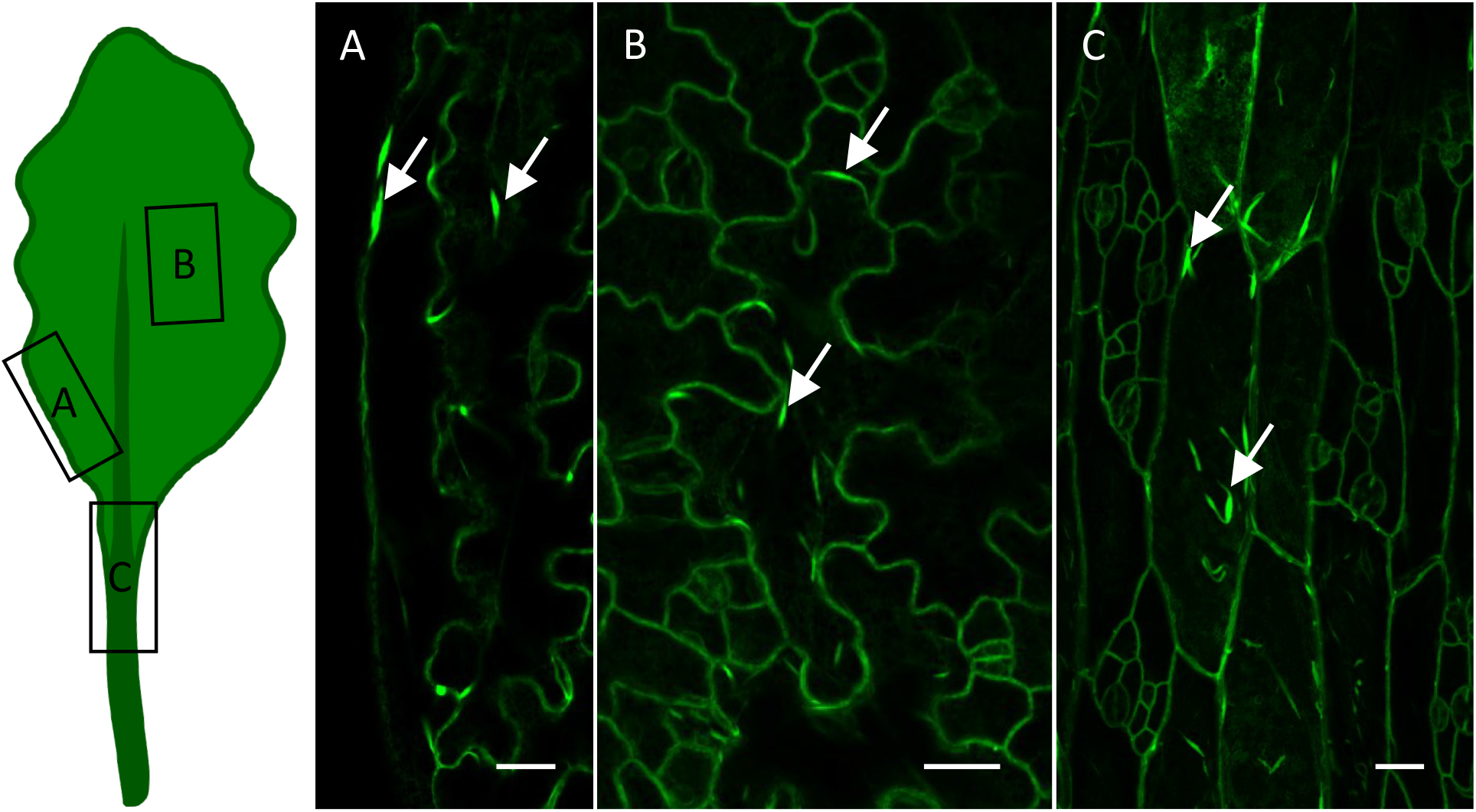
LER bodies in *Eutrema salsugineum* plants constitutively expressing ER-localized GFP-HDEL. Images showing LER body-containing giant cells (GCs) at **A**, the leaf margin, **B**, in the leaf blade, and **C**, in the petiole. Exemplary LER bodies are marked with arrows. Scale bars: 20 µm.

### LER bodies form during epidermal cell differentiation in *E. salsugineum*

To fully understand LER body formation during leaf development in *E. salsugineum*, an ∼2 mm long newly developing rosette leaf of a GFP-HDEL-expressing *E. salsugineum* plant was examined by confocal microscopy (Figure 5A). We assessed the development of rosette leaf epidermis by monitoring the cell shapes that were visualized by the cortical ER network at the surface of cells. The leaf base was characterized by small isodiametric cells, a typical characteristic of actively dividing cells, indicating that the cells are in the division phase or the so-called primary morphogenesis phase (Figure 5D). By contrast, the leaf tip contained much larger cells, which further differentiated into jigsaw puzzle-shaped epidermal cells, suggesting that these cells are in the so-called secondary morphogenesis phase, which includes cell expansion and differentiation. Therefore, the image shown in Figure 5A depicts epidermal cell differentiation and the switch from primary morphogenesis at the leaf base to secondary morphogenesis at the leaf tip. We closely examined different areas in the same leaf, and found that very small and roundish LER bodies had formed in small isodiametric cells during the primary morphogenesis phase at the leaf base (Figure 5D). However, only single cells accumulated LER bodies during this phase, indicating that these cells had already differentiated to establish a LER body-inducing gene expression profile. In the middle section of the leaf, the LER body-containing cells were identified as large pavement cells (Figure 5C). While the surrounding cells were still roundish and small, the LER body-containing cells had already started forming the lobe-neck shape of cell walls. This suggests the LER body-containing cells had stopped dividing via mitosis and had transitioned into the endoreduplication cycle, while the surrounding cells were still dividing. Fully developed cells could be recognized at the leaf tip, where only the large pavement cells contained LER bodies (Figure 5B).

**Figure 5:**
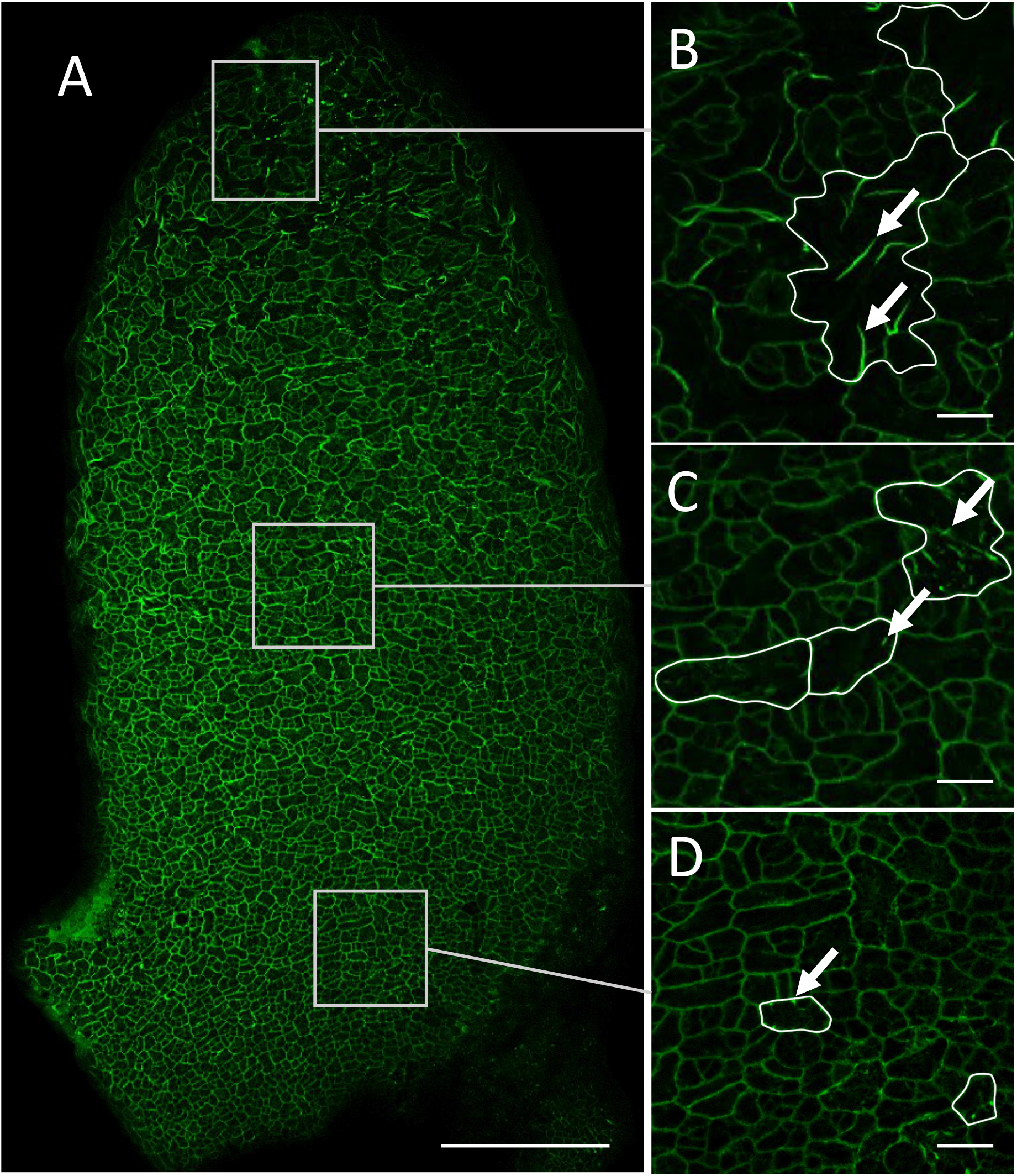
Large pavement cell differentiation in a newly emerging *E. salsugineum* rosette leaf. **A,** Confocal microscopy image of a newly emerging rosette leaf of a 25-d-old *E. salsugineum* plant expressing ER-localized GFP-HDEL (green). The development of large pavement cells containing ER bodies can be observed. **B-D**, Magnified images of the tip (B), middle section (C), and base (D) of the leaf showing the formation of LER bodies. Exemplary LER bodies in the developing large pavement cells are marked with arrows. Scale bars: 200 µm in A, 20 µm in B–D.

### ATML1 and LGO affect cER body-related gene expression to a lesser extent in cotyledons than in rosette leaves

Like the epidermal cells of rosette leaves, those of cotyledons are endoreduplicated (Moreno et al., 2020) and they contain many cER bodies (Matsushima et al., 2003b) in Arabidopsis. Therefore, we examined the expression levels of cER body-related genes (*NAI1*, *NAI2*, and *PYK10*) in 7-d-old seedlings of *ATML1*– and *LGO*– aberrant expression lines. The expression level of only *PYK10*, among all the tested genes, was significantly altered in *atml1*-*3* and *ATML1*-OX plants; *PYK10* expression was lower in *atml1*-*3* and higher in *ATML1*-OX compared with the WT (Figure 6A). The *LGO*-aberrant expression lines showed no significant change in the expression levels of *NAI1*, *NAI2*, and *PYK10*. Next, we determined NAI2 and PYK10 protein levels in the cotyledons of 7-d-old seedlings of *ATML1*– and *LGO*-aberrant expression lines (Figure 6B). Similar to the mRNA levels, protein levels barely changed in these lines, although the level of PYK10 was slightly lower in *atml1-3* and higher in *ATML1*-OX compared with the WT.

**Figure 6:**
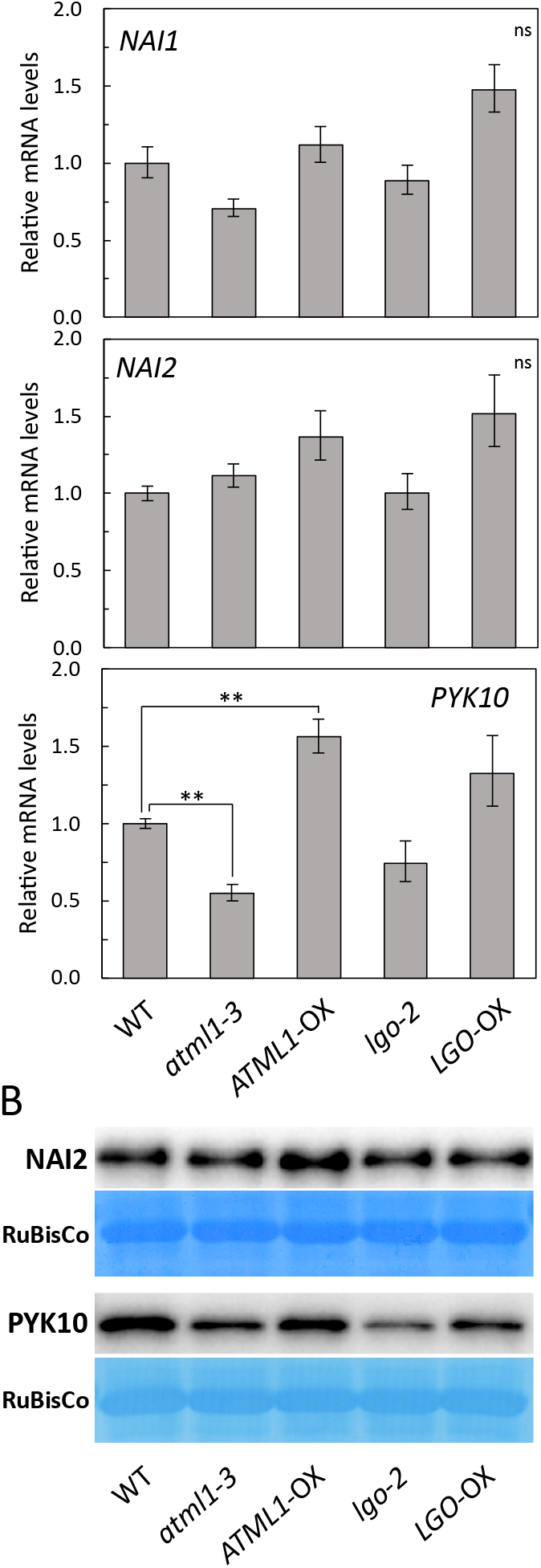
GC-aberrant Arabidopsis lines show only slightly altered levels of cER body-related mRNAs and proteins in cotyledons. **A,** Expression analysis of cER body-related genes in 7-d-old plants by qRT-PCR. Asterisks indicate significant differences compared with the WT (**p < 0.005; n = 3). **B,** Immunoblot analysis using anti-NAI2 and –PYK10 antibodies. *ATML1* as well as *LGO* slightly influenced the expression of cER body-related proteins.

### ATML1 and LGO greatly influence epidermal cell size distribution but only mildly influence cER body formation in cotyledons

We examined cER body formation on the adaxial side of the cotyledons of *ATML1*– and *LGO*-aberrant expression lines stably expressing GFP-HDEL, and then counted the cER bodies using a computational approach (Carpenter et al., 2006). Our results revealed a linear relationship between the number of ER bodies and the size of epidermal cells on the adaxial side of cotyledons in all genotypes (Figure 7A). The average cER body density in *ATML1*-OX (8.1 per 1000 µm^2^ leaf area) was similar to that in the WT (8.3 per 1000 µm² leaf area) but was reduced in *atml1*-*3* (6.9 per 1000 µm^2^ leaf area). Unexpectedly, cER body densities were reduced in both *lgo-2* and *LGO*– OX lines (5.8 and 4.9 per 1000 µm² leaf area, respectively). However, in contrast to the influence of *ATML1* on LER body formation in rosette leaves, we observed no substantial changes in cER body accumulation in the *ATML1*-aberrant expression lines. No cER bodies were observed in subsidiary cells located next to the guard cells in the WT, *ATML1*– and *LGO*-aberrant expression lines.

**Figure 7:**
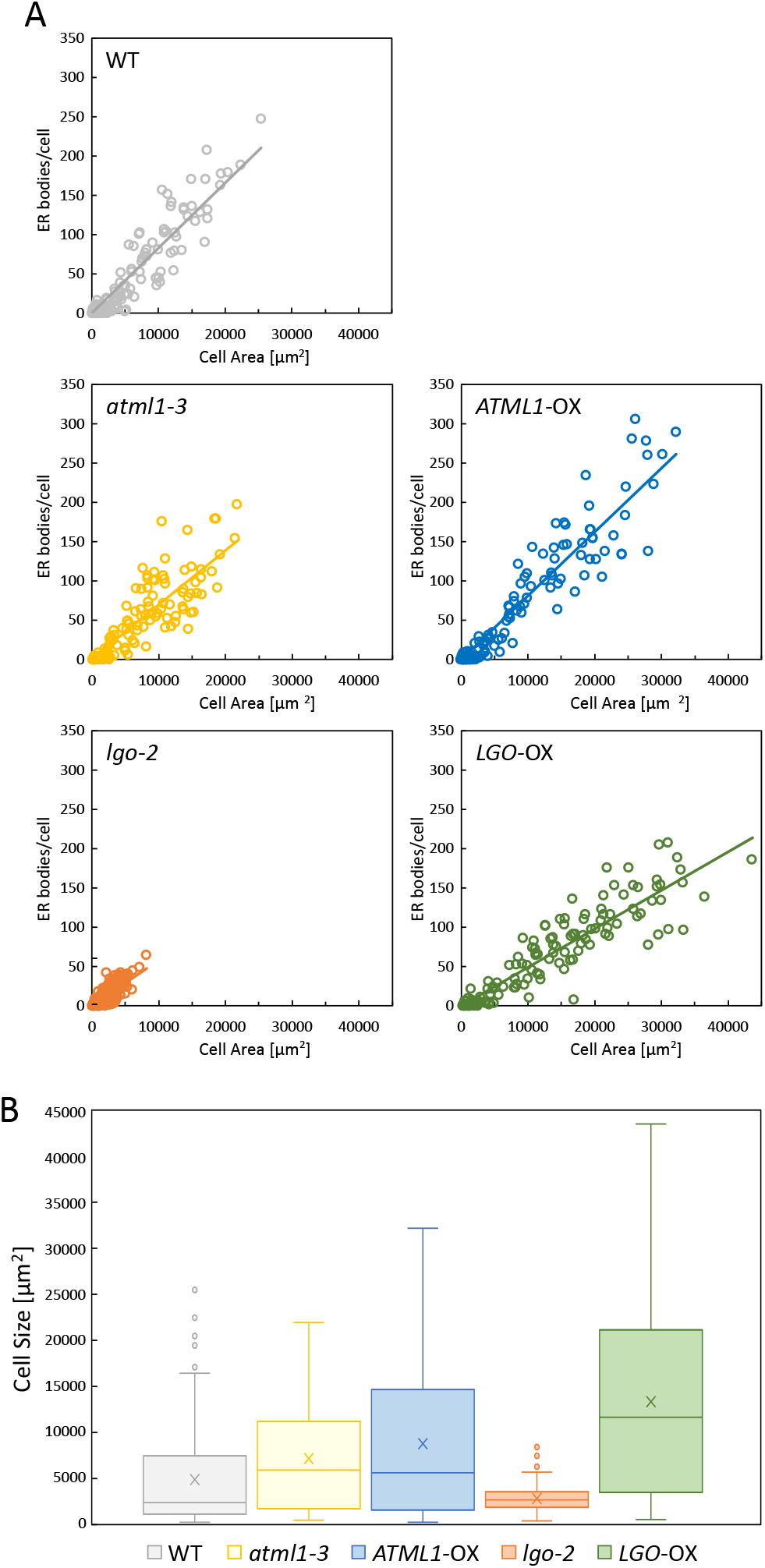
Influence of *ATML1* and *LGO* on cER body number and cell size in cotyledons. **A**, Analysis of the number of cER bodies per cell in the epidermal layer of WT, *atml1-3*, *ATML1*-OX, *lgo-2*, and *LGO*-OX cotyledons. The number of ER bodies was quantified relative to the area of each cell. Each circle represents a single analyzed cell. **B**, Epidermal cell area distribution in WT, *atml1-3*, *ATML1*-OX, *lgo-2*, and *LGO*-OX cotyledons. At least 127 cells were analyzed per genotype.

However, we observed a major difference in cell areas among the different genotypes (Figure 7B). Cells measuring up to 25,000 µm² were recorded in the WT. By contrast, cell areas were greatly reduced in *lgo*-*2* (8,000 µm² maximum) and considerably enhanced in *LGO-*OX (44,000 µm² maximum). The effect of *ATML1* expression on cell areas seemed to be lower than that of *LGO*; cell areas in the *ATML1-* OX line were enhanced to up to 32,000 µm², which is smaller than the maximum cell area observed in the *LGO*-OX line. Although the cell area range was slightly reduced in the *atml1*-*3* knockout line compared with the WT, the median value was 3-fold higher.

## Discussion

*ATML1* is a homeobox transcription factor responsible for epidermal cell differentiation (Iida and Takada, 2021). Our data indicate that ATML1 induces *BGLU18* as well as *NAI1*, whereby NAI1 is a direct regulator of *NAI2* and *PYK10* in Arabidopsis rosette leaves. Our findings suggest that, in addition to epidermal cell differentiation, ATML1 induces large pavement cell differentiation to equip a chemical defense based on the LER body system.

### *ATML1* induces LER body accumulation in large pavement cells in rosette leaves

In Arabidopsis rosette leaves, LER bodies are formed in large pavement cells, which are characterized by their physically large size, owing to endoreduplication (Melaragno, 1993). Because the properties of large pavement cells are similar to those of GCs in sepals, we hypothesized that the regulators of GC development also control large pavement cell differentiation. Indeed, our study indicates that the regulators of GCs, namely, *ATML1* and *LGO*, are involved in the formation of large pavement cells in rosette leaves. While the *ATML1*-aberrant expression lines showed considerable changes in the number of LER body-containing large pavement cells, the *LGO*-aberrant expression lines displayed changed epidermal cell sizes but negligible changes in LER body formation. These findings suggest that *ATML1* induces the functional differentiation of large pavement cells, and *LGO* promotes subsequent endoreduplication necessary for cell enlargement. Once the epidermal cells have differentiated into large pavement cells, these cells display a gene expression profile that enables LER body formation, regardless of their endoreduplication levels. Because ATML1 regulates both large pavement cell differentiation and GC differentiation, it is suggested that large pavement cells in rosette leaves and GCs in sepals constitute the same cell type, and ATML1 concomitantly promotes LER body-related gene expression. This hypothesis is indeed supported by the finding that several LER body-related genes exhibit differential expression between the sepals of GC-aberrant lines and those of the WT (Schwarz and Roeder, 2016).

In Arabidopsis sepals, *LGO* has been shown to exert a greater influence on GC– specific gene expression than *ATML1* (Schwarz and Roeder, 2016). Regarding LER body-related genes, *ATML1* had a stronger impact than *LGO*, although *LGO-*OX showed slightly enhanced expressions of LER body-related genes in rosette leaves. These data suggest that endoreduplication facilitates, but does not promote, the expression of LER body-related genes in the differentiated large pavement cells.

### *ATML1* is dispensable for cER body formation in cotyledons

Arabidopsis cotyledons are covered with endoreduplicated epidermal cells (De Veylder et al., 2002), which contain large numbers of cER bodies (Matsushima et al., 2003b). The size of cotyledonary epidermal cells was reduced in *lgo-2*, indicating that *LGO* is the major regulator of endoreduplication levels in the epidermal cells of Arabidopsis cotyledons. The number of cER bodies per cell increased linearly with the increase in cell area in WT plants and *ATML1*– and *LGO*-aberrant expression lines, indicating that cER body formation is directly correlated with cell size. Correlation between cell size and LER body formation was not observed in the rosette leaves of *LGO*-aberrant expression lines because *LGO*-OX produced large pavement cells without accumulating LER bodies, and *lgo-2* knockout mutants produced LER bodies in small pavement cells. Therefore, the results suggest that the regulatory mechanisms of cER bodies are different from those of LER bodies.

*ATML1* contributes to LER body formation and *NAI1* gene expression in rosette leaves, but is dispensable for cER body formation in cotyledonary epidermal cells. *ATML1* overexpression enhanced epidermal cell areas, although not as much as *LGO* overexpression, and the knockout mutation of *ATML1* did not decrease cell areas. These findings indicate that ATML1 exhibits low-level activity in cotyledons, and suggests that an alternative transcription factor regulates *LGO* and possibly cER body–related genes. One candidate transcription factor-encoding gene is *PROTODERMAL FACTOR 2* (*PDF2*), which is highly homologous to *ATML1* in Arabidopsis. *PDF2* and *ATML1* are known to act redundantly (Ogawa et al., 2014). Therefore, *PDF2* may be responsible for rescuing *ATML1* deficiency in the *atml1*-*3* knockout line. Consistent with this finding, the double knockout mutation of Arabidopsis *ATML1* and *PDF2* is lethal because of the lack of epidermal cell differentiation (Abe et al., 2003), indicating that these homeobox genes redundantly regulate epidermal cell differentiation, *LGO*-mediated cell expansion, and possibly cER body formation.

### Regulatory pathway of LER body formation

We found that large pavement cell differentiation and LER body formation are co-regulated by ATML1. ATML1 regulates LER body-related genes, including *NAI1*, *NAI2*, *PYK10*, and *BGLU18*. It is unlikely that ATML1 directly regulates *NAI2* and *PYK10* expression, since the knockout mutation of *NAI1* is enough to suppress PYK10 and NAI2 protein translation in large pavement cells in rosette leaves (Nakazaki et al., 2019a). Additionally, the overexpression of *NAI2* and *PYK10* in *ATML1*-OX was abolished after introduction of the *nai1*-*1* mutation, supporting the theory that ATML1 indirectly regulates the expression of *NAI2* and *PYK10* (Figure 3). Therefore, we propose a regulatory scheme in which ATML1 directly regulates the expression of *NAI1* and *BGLU18*, and ATML1-induced NAI1 regulates the expression of *NAI2* and *PYK10* (Figure 8). Consistently, we found L1 boxes in the promoter regions of *NAI1* and *BGLU18*, to which ATML1 can bind to activate gene transcription (Abe et al., 2001). We could not find L1 boxes in the promoter regions of *NAI2* and *PYK10*, but NAI1 has been shown to bind to the promoter regions of these genes to upregulate their expression (Sarkar et al., 2020), indicating that NAI1 directly regulates *NAI2* and *PYK10* expression during LER body formation. We also found LER bodies in *E. salsugineum* as well as a potential ATML1-binding site in the promoter region of the *E. salsugineum NAI1* homolog. This suggests that the regulatory pathway controlling ATML1-mediated LER body formation is conserved among Brassicaceae family plants.

**Figure 8:**
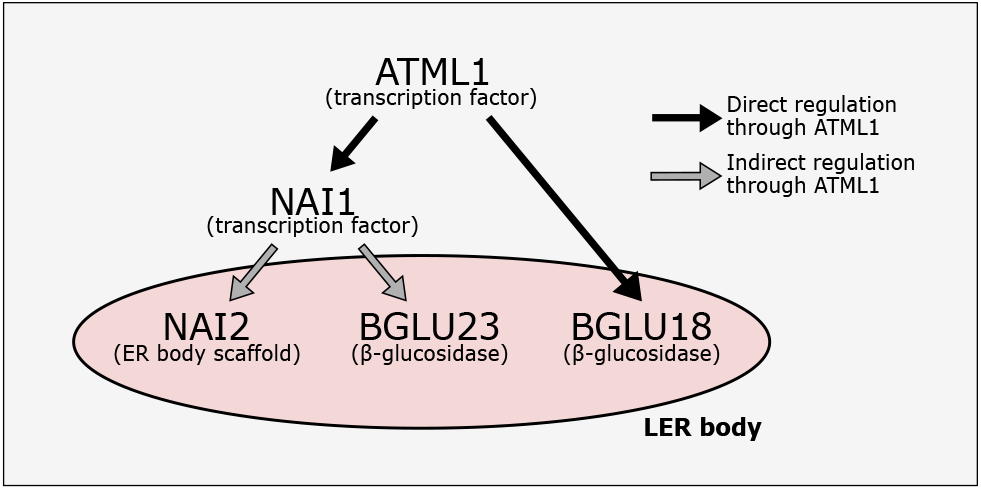
A proposed gene regulatory scheme for the formation of LER bodies in Arabidopsis.

Although *LGO* did not act as a primary regulator of LER body formation, it facilitated the expression of LER body-related genes, suggesting that endoreduplication influences gene expression. Recent findings suggest that *LGO* affects gene expression levels (Schwarz and Roeder, 2016), and ATML1 functions in a dose-dependent manner to regulate GC differentiation in Arabidopsis sepals (Meyer et al., 2017). Together with these findings, our results imply that LGO enhances LER body-related gene expression by increasing gene dosage in large pavement cells in rosette leaves. Moreover, the JA receptor COI1 has been shown to regulate *BGLU18* expression in rosette leaves (Nakazaki et al., 2019a, 2019b). Although COI1 is not involved in LER body formation (Nakazaki et al., 2019a, 2019b), the findings of this study and previous studies suggest the co-regulation of *BGLU18* by *COI1* and *ATML1* in rosette leaves. Further investigation is needed to clarify the role of COI1 in LER body formation. Overall, our findings highlight the unique function of ATML1 in protecting plants from herbivores; ATML1 is not only responsible for the differentiation of the physical outer barrier of plants, the epidermis (Iida and Takada, 2021), but also for the formation of a chemical barrier in the form of LER bodies (Nakazaki et al., 2019a; Roeder et al., 2012; Nakazaki et al., 2019b).

## Methods

### Plant material and growth conditions

*Arabidopsis thaliana* genotypes used in this study are described in Supplemental Table 2. Arabidopsis plants were grown for 14 days on half-strength Murashige and Skoog (½ MS) solid medium (1% [w/v] sucrose, 0.5% [w/v] MES-KOH buffer [pH 5.7], 0.4% [w/v] gellan gum, and 0.5× MS salts [Wako]) under sterile controlled conditions (22

°C, continuous light, and ∼100 mE s^-1^ m^-2^ light intensity). Subsequently, the plants were transferred to Jiffy peat (41 mm) and grown under a 16 h light/8 h dark cycle. *Eutrema salsugineum* plants were grown from seeds (NASC ID: N22504) under non-sterile controlled conditions (20 °C and 16 h light/8 h dark cycle).

### Plant transformation

*Agrobacterium tumefaciens* strain C58C1Rif containing the binary vector pBI121, which harbored the DNA sequence encoding signal peptide-GFP-His-Asp-Glu-Leu (SP-GFP-HDEL) fusion under the control of the constitutive 35S promoter (Yamada et al., 2008) was used for stable transformation of *atml1*-3, *ATML1*-OX, *lgo*-2 and *LGO*-OX Arabidopsis lines as well as *E. salsugineum* plants. Transformation was performed *via* floral dipping as described previously (Zhang et al., 2006). To transform *E. salsugineum*, the dipping procedure was repeated by using the same plants once a week over a period of 4 weeks.

### Production of ATML1 OX/nai1-1 plants

*ATML1* OX plants with *nai1*-*1* background were produced by crossing ATML1 OX and *nai1*-*1* plants. To ensure homozygosity of the *nai1*-*1* point mutation, the *NAI1* region covering the point mutation site was amplified from several F3 generation plants using the ‘Phire Plant Direct PCR Master Mix’ (Thermo Fisher). Presence of the homozygote point mutation was verified by Sanger sequencing. *ATML1* OX causes an elongated rosette leave phenotype, therefore homozygosity of *ATML1* OX was ensured by observing the plant phenotypes of the F4 generation.

### Gene expression analysis

Total RNA was extracted from either 7-d-old cotyledons (12 cotyledons from six plants per sample, with three replicates per genotype) or 14-d-old plants with roots and cotyledons removed (three plants per sample, with three replicates per genotype) using TRIzol (Thermo Fisher Scientific), according to the manufacturer’s instructions. The isolated total RNA was resuspended in 100 µL of water. To remove possible genomic DNA (gDNA) contamination, 1 µg of each isolated RNA sample was treated with DNase I (Sigma Aldrich). The DNase I-treated RNAs were reverse transcribed using Ready-to-Go RT-PCR beads (GE Healthcare Life Science) and random oligomers, and the obtained cDNA samples were stored at –20 °C. Real-time PCR was performed using PowerUp SYBR Green Master Mix (Thermo Fisher Scientific), according to the manufacturer’s instructions, and QuantStudio 6 (Thermo Fisher Scientific). Gene-specific primer sets (Supplemental Table 3) were designed using Primer3Plus (http://primer3plus.com) to produce products 80–150 bp in length. *UBQ10* was used as an internal control gene.

### Immunoblot analysis

Protein extraction was performed using either 7-d-old cotyledons (48 cotyledons from 24 plants per sample, with three replicates per genotype) or the first pair of rosette leaves collected from 14-d-old plants (eight leaves collected from four plants per sample, with three replicates per genotype). To extract total protein, the plant material was weighed and treated with 8-fold excess of sample buffer (8% [w/v] sodium dodecyl sulfate [SDS], 0.1 M Tris-HCl [pH 6.8], 20% [v/v] glycerol, 2% [v/v] saturated bromophenol blue, and 12% [v/v] 2-mercaptoethanol).

To detect proteins in cotyledons, each 3 µL of isolated protein extracts was applied to 10% SDS gel. The amount of protein extract that was loaded on the gels varied depending on the primary antibody (4, 5, and 8 µL for anti-NAI2, –BGLU18, and – PYK10 antibodies, respectively). Immunoblotting was performed as described previously (Yamada et al., 2008). PYK10 was detected using Can Get Signal (TOYOBO) (1:10,000 dilution). RuBisCo (loading control) was visualized by staining with Coomassie Brilliant Blue.

### Microscopy

Fluorescent proteins and dyes were detected using a confocal laser scanning microscope (LSM810; Carl Zeiss). To visualize the cell wall in Arabidopsis rosette leaves, the leaves were incubated in propidium iodide (PI) staining solution (100 µg/mL) for 15 min on ice; the incubation was performed on ice to reduce ER body movement to a minimum. Depending on the experiment, z-stacks were captured with a slice thickness of 3 µm. GFP signal was detected using an Argon laser (488 nm), and PI staining was visualized using a laser with a wavelength of 561 nm. All pictures were taken from the adaxial side of the leaf.

### Determination of petiole coverage with LER body-containing cells

To quantitively measure the differences in LER body formation among *atml1*-*3*, *ATML1*-OX, *lgo*-*2*, and *LGO*-OX relative to the WT, five z-stack images were captured from the epidermal cell layer of each of the GFP-HDEL-expressing lines. Cell boundaries were visualized by incubating the leaves in PI, as described above. Both green (ER) and red (PI) signals were detected. Images of the petiole area adjacent to the leaf blade were captured. The z-stacks of each image were merged using maximum intensity projection, and cell boundaries were overlaid manually using Inkscape, a free vector graphics software. Areas of cells containing LER bodies as well as those devoid of LER bodies were then measured using FIJI software. Stomatal cells were not taken into consideration and were not included in the area of cells lacking LER bodies.

### Correlation between cER body number and cell size

CellProfiler (Carpenter et al., 2006) was used to analyze the number of cER bodies per unit area and to determine whether the number of cER bodies per cell depends on the cell size or on the ploidy level of the individual cell. The PI-stained cotyledons of WT, *atml1*-*3*, *ATML1*-OX, *lgo*-*2*, and *LGO*-OX plants expressing GFP-HDEL were analyzed. The z-stack images of 9-d-old cotyledons were merged using maximum intensity projection. ER bodies and cell boundaries were automatically detected via the green and red channels, respectively, and the cell boundaries then manually edited. Next, the ER bodies were assigned to the cells and the cell area of each cell and the corresponding number of ER bodies was measured and alaysed.

### Ultra-performance liquid chromatography-tandem mass spectrometry (UPLC-MS/MS) analysis

Collected frozen leaves (∼200 mg) were homogenized in DMSO (2.5 μl per 1 mg fresh weight), and centrifuged. The supernatant was collected and analyzed using Acquity UPLC system (Waters, USA) coupled with a micrOTOF-Q mass spectrometer (Bruker Daltonics, Germany) as described earlier (Czerniawski et al., 2021). The obtained raw data were processed by MZmine v2.32 software (Pluskal et al., 2010) including ion detection and chromatogram building. Subsequently, the extracted ion chromatograms were deconvoluted, aligned and gap-filled. Finally, the data matrix was limited to chromatograms of molecular ions corresponding to glucosinolates identified as described earlier (Czerniawski et al., 2021). The amounts of particular glucosinolates were estimated by calculation of respective peak areas in molecular ion chromatograms.

### Statistical analysis

In the qPCR analysis and the comparison of petiole areas covered with ER body-containing cells in different genotypes, Student’s t test was used for pairwise comparison of each mutant and wild-type plants. Glucosinolate contents were analysed with ANOVA followed by Tukey HSD, whereby significant changes compared to WT were indicated in the graph.

### Accession numbers

Sequence data from this article can be found in the Arabidopsis Genome Initiative (AGI) or GenBank/EMBL databases under the following accession numbers: At4g21750 (ATML1), At3g10525 (LGO), At2g22770 (NAI1), At3g15950 (NAI2), At3g09260 (PYK10), At1g52400 (BGLU18), and LOC18021790, XM_006404677 (*E*.

*salsugineum NAI1* homolog). The accession numbers of the T-DNA insertion mutants are listed in Supplemental Table 2.

## Supplemental data

**Supplemental Table 1:** Normalized peak areas of individual glucosinolates measured during LC/MS analysis of samples obtained from leaves of the indicated genotypes.

**Supplemental Table 2:** Overview of *Arabidopsis thaliana* genotypes employed in this study.

**Supplemental Table 3:** List of qRT-PCR primers used in the study.

**Supplemental Figure 1:** Relative glucosinolate contents of the first leaf pair of 14-d-old *ATML1-*and *LGO-* aberrant expression Arabidopsis lines.

**Supplemental Figure 2:** Schematic representations of the promoter regions of LER body-related Arabidopsis *NAI1*, *BGLU18*, *NAI2*, and *PYK10* genes and the *E. salsugineum NAI1* homolog (LOC18021790).

**Supplemental Figure 3:** Influence of *ATML1* on *BGLU18* mRNA and protein levels in 7-d-old cotyledons.

## Acknowledgments

We thank Adrienne H.K. Roeder (Cornell University) and Gwyneth C. Ingram (Université de Lyon) for sharing the Arabidopsis *ATML1*-OX seeds. This work was supported by the National Science Centre of Poland (OPUS grant UMO-2020/37/B/NZ3/04176 to K.Y., and PRELUDIUM grant UMO-2020/37/N/NZ3/03591 to A.W.), and by institutional support provided by the Malopolska Centre of Biotechnology, Jagiellonian University. The open-access publication of this article was funded by the BioS Priority Research Area under the program Excellence Initiative – Research University at the Jagiellonian University in Krakow.

## Author contributions

A.W. and K.Y. designed the research. A.W. designed and performed the experiments and analyzed the data. A.W. generated transgenic plant lines. P.C. and P.B. performed glucosinolate analyses. A.W., M.L.-K., P.B. and K.Y. wrote the manuscript.

